# Biomimetic virus-like mesoporous silica nanoparticles activate NK cells indirectly via monocyte crosstalk

**DOI:** 10.64898/2026.04.22.720074

**Authors:** Minna Sivonen, Silja Saarela, Jiajia Wang, Mira Saari, Emmi Järvelä, Lotta Andersson, Enkhzaya Batnasan, Leena Latonen, Helka Göös, Vesa-Pekka Lehto, Wujun Xu

## Abstract

Cancer immunotherapies show clinical promise but often rely on T-cell priming and are limited by tumor heterogeneity and the immunosuppressive tumor microenvironment (TME). Innate immune activation offers a complementary strategy, with specific aim in natural killer (NK) cell activation for antigen-independent response. Biomimetic nanoparticles combining virus-like morphology with cell membrane (CM) coating offer a strategy to engage this innate immune axis. This study investigates virus-like mesoporous silica nanoparticles (VLPSi) with tunable spikes, surface functionalization, and CM coating as innate immunity modulators. Optimization revealed that longer spikes, amine functionalization, and CM coating synergistically enhance NK cell activation within human PBMCs, as indicated by CD69/CD25 upregulation and IFN-γ secretion. CD14^+^ monocyte depletion attenuated activation, identifying monocyte-dependent crosstalk as a key mechanism. In purified NK cells, engineered CM-coated VLPSi induced early activation and supported feeder-free expansion. These results define topology, surface chemistry, and CM coating as parameters for innate immune modulation.

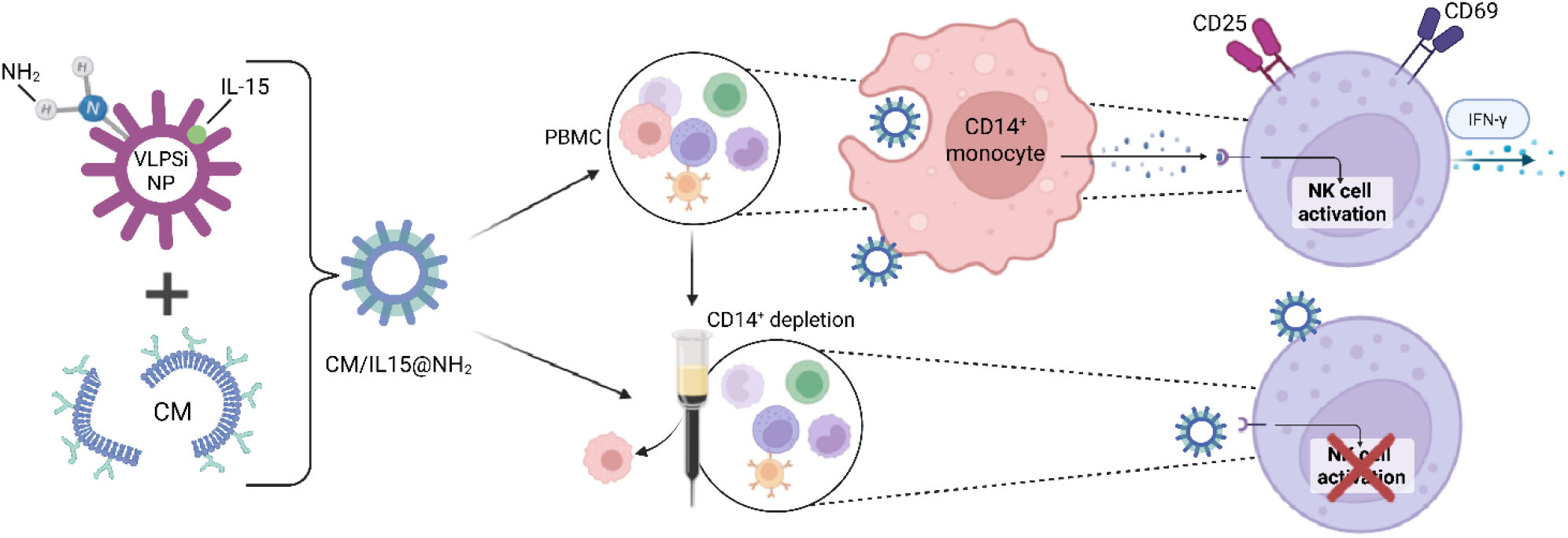

## Introduction

Recent advances in cancer immunotherapy have transformed the therapeutic landscape. Approaches such as immune checkpoint inhibitors [1], adoptive cell therapies [2], and cancer vaccines [3,4] harness the patients’ immune system to eliminate malignant cells. However, many of these strategies rely on antigen-specific T-cell priming and cannot overcome tumor heterogeneity and patient variability, resulting in modest response rates and short-term therapeutic benefits [5]. Therapies that activate the innate immune response, particularly natural killer (NK) cells, do not rely on recognition of a single antigen and therefore hold potential for broader antitumor responses [6]. NK cells have demonstrated efficacy against hematological tumors, but their effector function and infiltration in solid tumors are often suppressed by the immunosuppressive tumor microenvironment (TME). In addition, interactions with other immune cells have an impact on their activation. For example, macrophages prime NK cells via cytokine signaling (*e*.*g*. IL-12, IL-18) and direct cell-cell contact [7]. This crosstalk represents an important aspect of innate immune activation and offers mechanisms that could be used in nanomaterial design for cancer immunotherapies.

Nanoparticle (NP) designs that combine activating ligands and cytokine presentation could offer a strategy to mimic NK cell-activating signals and trigger robust innate immune responses. NPs engineered with biomimetic features, including virus-like surface morphology and biological components such as CM coating, offer a versatile platform to integrate synthetic nanomaterials with biologically relevant cues for cancer immunotherapies [8,9]. Virus-like surface topology can enhance immune activation by mimicking pathogen-associated structural cues, promoting uptake by antigen-presenting cells (APCs) and stimulating innate immune responses [10]. In parallel, CM-coating transfers surface proteins, adhesion molecules, and immunologically relevant ligands from their parent cells onto the NPs [11,12]. Depending on the parent cell, CM coating can be utilized for homotypic targeting, prolonged circulation and improved biocompatibility [13]. CMs derived from cancer cells have shown potential to influence NK cell responses through cancer-specific antigen presentation [8]. Although virus-like nanostructures and CM-coated spherical nanoparticles have independently demonstrated promise for cancer immunotherapy, their combined effects remain largely underexplored. Given that innate immune response relies on rapid, coordinated communication between myeloid cells and NK cells, cancer immunotherapies capable of engaging this axis could offer significant advantages. This represents an opportunity to design NPs that can more effectively modulate innate immune response.

In this study, virus-like mesoporous silica nanoparticles (VLPSi) were engineered with rigid tunable spikes and biomimetic cancer CM coating. Mesoporous silica is widely used for biomedical applications due to its biocompatibility, structural stability, and high drug-loading capacity [14]. Advances in nanotechnology enable precise control over its particle size, surface morphology, and chemistry [15,16]. Virus-like surface topology is known to enhance cellular uptake and can stimulate innate immune responses by mimicking pathogen-associated structural cues [17,18]. Although direct evidence for NK cell activation by virus-like NPs is limited, these cues can engage in innate immune pathways that support NK cell function [19]. We aimed to investigate how VLPSi morphology, surface functionalization, CM source, and cytokine incorporation influence NK cell activation within human peripheral blood mononuclear cells (PBMCs). These results reveal how NP design features can be utilized to harness innate immune interactions to drive NK cell activation, establishing design principles for innate immune engaging NPs for next-generation cancer immunotherapies.

## Results

VLPSi were engineered using a single-micelle epitaxial growth approach in a biphasic reaction system where hexadecyltrimethylammonium bromide (CTAB) served as a template and tetraethyl orthosilicate (TEOS) as a precursor. During synthesis, virus-like spikes form through the assembly of CTAB and silica oligomers in the presence of cyclohexane. By modulating reaction time (48 to 72 hours), VLPSi with approximately 5 nm (VLPSi-5) and 30 nm (VLPSi-30) virus-like spikes were synthesized. VLPSi with 30 nm spikes have previously demonstrated higher cellular internalization and immune engagement [19], and were selected as the primary platform for subsequent studies, while VLPSi-5 were used as a reference control (Supplementary Figure S2A).

The influence of spike length and CM coating to cellular uptake was assessed using FITC-labelled VLPSi-5 and VLPSi-30. CMs derived from IFN-γ treated MDA-MB-231 (MDA) cells (γCM) served as a source of tumor-associated antigens and biologically relevant ligands. VLPSi and γCM-coated VLPSi (γCM@VLPSi) were co-cultured with human breast cancer cells (MDA), monocytes (THP-1) and expanded primary human NK cells (exNK). Flow cytometry results revealed that the uptake profile was cell-type dependent and enhanced by both γCM coating and longer spikes, with γCM@VLPSi-30 showing the highest uptake (Figure 1A). MDA cells had the highest uptake profile, followed by THP-1, while exNK cells showed minimal uptake, with modest increase observed for γCM@VLPSi-30. Based on these results, VLPSi-30 were selected for subsequent studies.

**Figure 1.**
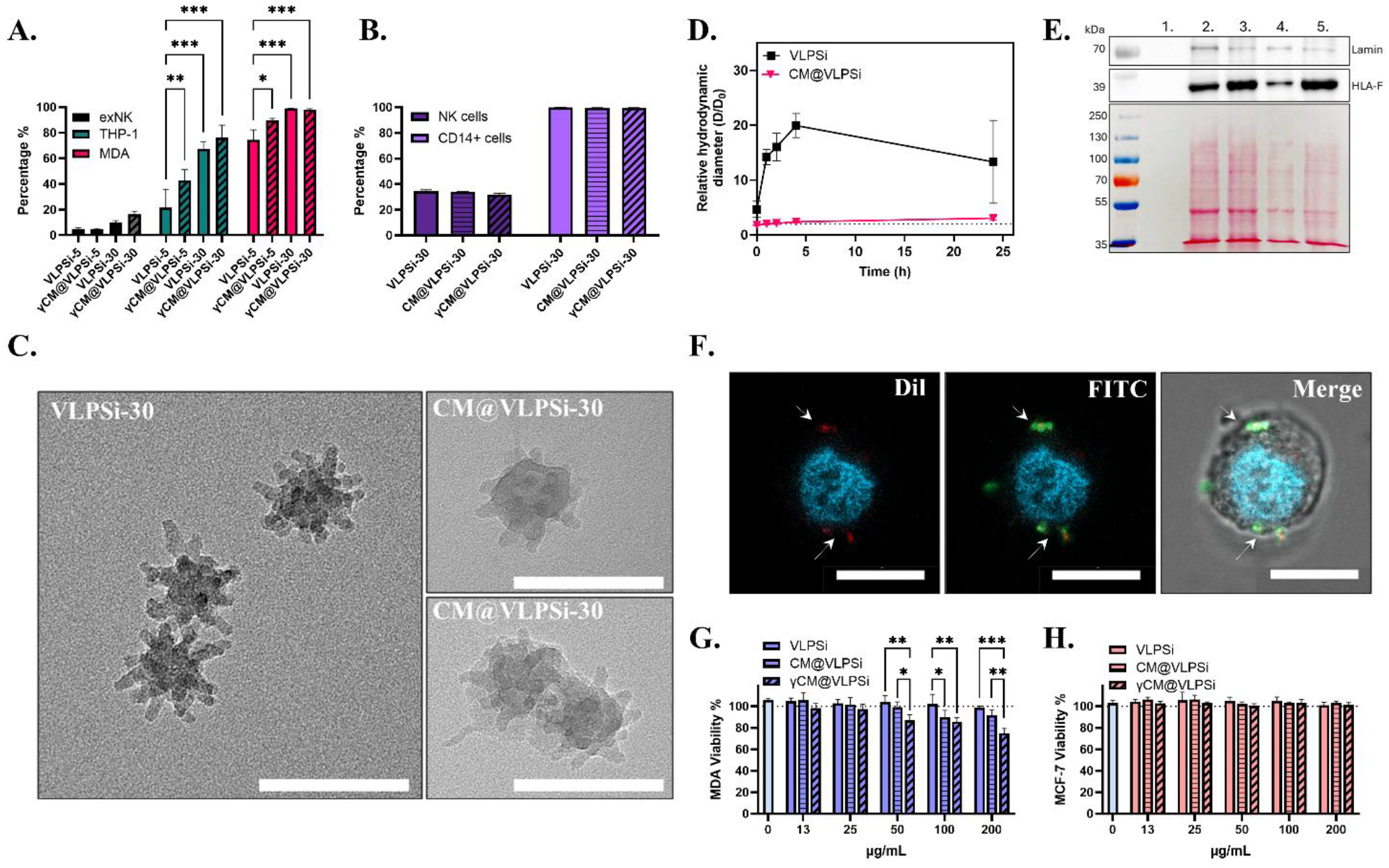
Characterization of virus-like mesoporous silica nanoparticles (VLPSi). **A.**Bar graph showing the percentage of FITC positive expanded NK cells (exNK), THP-1 and MDA-MB-231 (MDA) cells after 4-hour co-culture with FITC labelled VLPSi carrying 5 nm (VLPSi-5) or 30 nm (VLPSi-30) spikes, with and without IFN-γ treated MDA cell membrane (γCM) coating (γCM@VLPSi-5, γCM@VLPSi-30). **B**. Bar graph showing the percentage of FITC positive naïve NK cells and CD14^+^ cells within PBMC after 4-hour co-culture with FITC labelled VLPSi-30 with and without CM coating. CMs were derived from MDA (CM; striped), and IFN-γ stimulated MDA (γCM; diagonal pattern). **C**. Transmission electron microscope (TEM) images from VLPSi with 30 nm spikes (VLPSi-30), and MDA CM-coated VLPSi (CM@VLPSi-30). **D**. Relative hydrodynamic diameter (D/D_0_) of VLPSi and CM-coated VLPSi (CM@VLPSi) in PBS in 1, 2, 4 and 24 hour timepoints. **E**. Western blotting analysis for HLA-F (MHC-I) marker, Lamin B as a loading marker. SDS-PAGE showing protein bands of (1.) bare VLPSi-30, CM lysates from (2.) MDA and (3.) IFN-γ stimulated MDA (γCM), and VLPSi-30 coated with (4.) MDA-CM and γCM (5.). Samples were run at equal protein concentration (25 µg) and stained with Ponceau S. **F**. Representative confocal image showing the co-localization (white arrows) of VLPSi (green) and CM (red) upon cellular uptake in THP-1 cells. **G-H**. Bar graphs showing the biocompatibility (viability) of **G**. MDA and **H**. MCF-7 cell lines after 24 hours co-culturing with 13, 25, 50, 100, and 200 µg/mL of VLPSi-30 with and without CM and γCM coatings. Data shown as mean ±SD. ***p < 0.001, **p < 0.01, *p < 0.05. Scale bars: TEM (A), 100 nm; confocal images (E), 10 µm. Data shown as mean ±SD, n=3.

To further investigate the distribution within a heterogenous immune cell population, VLPSi-30 were co-cultured with PBMC, and the impact of CM coatings was assessed using CMs derived from MDA (CM) and IFN-γ treated MDA cells (γCM). In this context, CD14^+^ monocytes dominated the uptake regardless of CM source, whereas NK cells consistently showed lower uptake (Figure 1B). This observation goes together with the known phagocytic capacity of monocytes and indicates that VLPSi primarily interact with myeloid cells within PBMCs.

Transmission electron microscopy (TEM) characterization of VLPSi-30 shows their virus-like morphology, featuring surface spikes and an overall diameter of approximately 80 nm (Figure 1A; VLPSi-30). TEM images of CM-coated VLPSi-30 (CM@VLPSi) confirmed the presence of a CM surrounding the particles, indicating successful membrane coating (Fig. 1C; free CM shown in Supplementary Fig. S2B). CM coating improved colloidal stability in PBS, compared to bare VLPSi-30 (Figure 1D). Bare VLPSi-30 aggregated rapidly, whereas the diameter of CM@VLPSi remained stable over 24 hours, indicating suitability for biological applications. Preservation of membrane proteins during coating process was confirmed using SDS-PAGE and Western blotting. IFN-γ-treated MDA were included to the study to modulate MHC-I expression on MDA cells and assess its impact on NK cell activation. Results show that CM and γCM-coated VLPSi-30 preserved their distinct protein bands during the CM coating process, including IFN-γ-associated upregulation of MHC-I marker (HLA-F), seen in free γCM and γCM-coated VLPSi samples (lanes 3 and 5; Figure 1E). The co-localization of VLPSi (green) and CM (red) upon cellular uptake was confirmed using confocal imaging (Fig. 1F).

Biocompatibility was assessed using MDA and MCF-7 breast cancer cells. Cells were co-cultured with bare VLPSi-30 and CM-coated VLPSi-30 (CM@VLPSi-30, and γCM@VLPSi-30) in a concentration dependent manner for 24 hours. Bare VLPSi-30 did not show any detectable impact on cell viability even at highest concentration of 200 µg/mL (Figure 1F and 1G), consistent with the known safety profile of mesoporous silica [22]. In contrast, CM@VLPSi-30 modestly reduced viability in MDA cells (Figure 1F), reflecting the homotypic CM targeting, while γCM@VLPSi-30 caused 20% decrease in cell viability at highest concentrations compared to bare VLPSi. These results suggest that IFN-γ induced alterations in CM composition could enhance homotypic interactions without causing off-target toxicity. Together, these structural and biochemical characteristics confirmed the presence of CM on VLPSi, supporting their use in subsequent immunotherapy applications.

NK cell activation requires stimulatory signals, such as IL-15, that are often absent in resting PBMCs. To evaluate the ability of VLPSi-30 to modulate NK cell activation within a complex immune population, PBMCs were co-cultured with VLPSi-30 in the presence or absence of interleukin-15 (IL-15), a cytokine known to support NK cell proliferation and functional activation. NK cell activation was assessed by CD69 and CD25 expression using flow cytometry. In physiological conditions, monocytes prime NK cells through co-stimulatory interactions and cytokine signaling, with IL-2, IL-15, and IL-21 being the key mediators for NK cell activation and proliferation [25]. However, systemic administration of these cytokines is often limited by their off-target toxicity [26]. Nanoparticle based strategies could offer a solution for more localized delivery strategies. Previous studies have demonstrated that IL-15 integrated nanoparticles can enhance IFN-γ production and NK cell infiltration at tumor site [27]. This is consistent with our findings. In the absence of IL-15, CM-coated VLPSi-30 (CM@VLPSi-30) induced only a modest, increase in CD69^+^ NK cell population compared to bare VLPSi and medium controls, whereas CD25^+^ NK cell population remained low (Fig. 2A), indicating limited activation without IL-15. When IL-15 (10 ng/mL) was supplemented to the medium (+ IL-15) or integrated onto the VLPSi-30 (IL15@VLPSi-30), both CD69 and CD25 expressions were upregulated across all groups and further increased with CM coating (Fig. 2B). These results demonstrate that IL-15 is essential for robust NK cell activation and that VLPSi can serve as a platform for co-delivery of cytokines and CM ligands.

**Figure 2.**
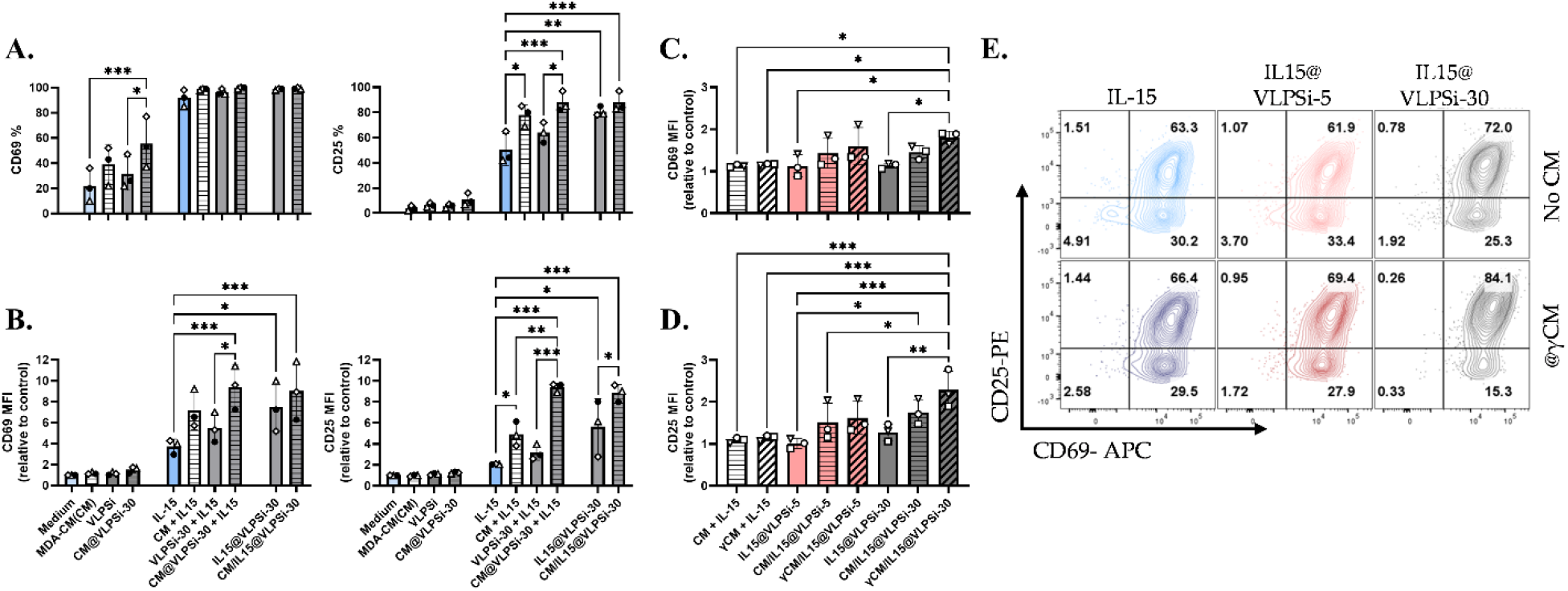
IL-15 and virus-like morphology’s impact on NK cell activation. **A-B.** Bar graphs show the percentage of **A**. CD25 and **B**. CD69 positive NK cells and **B**. Median Fluorescence Intensity (MFI) of CD69 and CD25 expression on NK cells normalized to medium control. Peripheral blood mononuclear cells (PBMCs) were co-cultured with virus-like mesoporous silica nanoparticles (VLPSi) with 30 nm spikes (VLPSi-30; grey) and MDA-MB-231 cell membrane (CM)-coated VLPSi (CM@VLPSi-30; striped) for 24-hours with and without 10 ng/mL IL-15 in the media or integrated with VLPSi (IL15@VLPSi). IL-15 (blue) and CM alone (white) were used as controls. **C-D**. Bar graphs comparing VLPSi with 5 nm spikes (VLPSi-5;pink) and 30 nm spikes (VLPSi-30; grey) impact to **C**. CD69 and **D**. CD24 expression with and without CM coating (CM from untreated MDA (CM; striped) CM or IFN-γ treated MDA CM (γCM; diagonal pattern)) after 24 hour co-culturing. Free CMs (white) used as controls. **E**. Representative contour plot of CD25 and CD69 co-expression on NK cells. Data shown as mean ±SD. ***p < 0.001, **p < 0.01, *p < 0.05. Donors represented by different symbols, n=3.

After establishing that IL-15 is required for robust NK cell activation, the impact of MHC-I upregulation and spike length were studied. IL-15-integrated VLPSi-30 (IL15@VLPSi-30) with 5 nm (VLPSi-5) and 30 nm (VLPSi-30) spikes were coated with CM derived from untreated (CM) or IFN-γ stimulated MDA cells (γCM). As previously shown (Figure 1C), γCM has higher MHC-I expressions than standard MDA CM. Across all donors, γCM coating consistently induced strongest NK cell activation. γCM/IL15@VLPSi-30 significantly upregulated CD25 expression compared to bare IL15@VLPSi-5 and IL15@VLPSi-30, and free CM or γCM controls (Figure 2D). CD69 expression followed a similar trend, with γCM/IL15@VLPSi-30 inducing the highest expression (Figure 2C). Co-expression analysis revealed that VLPSi with 30 nm spikes induced the largest population of CD69^+^ CD25^+^ NK cells, reaching 84% double-positive cells (Figure 2E). Although higher MHC-I expression is typically associated with NK cell inhibition, γCM coating promoted activation. This suggests that other stimulatory factors present in γCM, could override MHC-I-mediated inhibition. IFN-γ treatment has been shown to induce the expression of NK cell activation ligands, such as ICAM-1 on MDA-MB-231 cells [28], which could outweigh the MHC-I mediated inhibition.

These results indicate that NK cell activation is influenced by cytokine delivery, CM coating, and spike length, with 30 nm spikes producing the strongest activation. Therefore, VLPSi-30 was selected for all subsequent studies as the most effective platform for NK cell stimulation.

Having established that VLPSi with longer surface spikes (30 nm) enhance NK cell activation, the influence of surface chemistry was evaluated using baseline MDA CM coating. VLPSi-30 were functionalized with amino (NH_2_), hydroxylamine (NOH), or quaternary amine (NCH_3_) to investigate how surface chemistry and charge influence NK cell activation. Surface functionalization and membrane coating were assessed by changes in zeta potential (ζ). Bare VLPSi-30 had a negative charge, which shifted to positive after amino modifications and was subsequently restored to a level comparable to free CM after CM coating (Figure 3A), indicating a successful modification and presence of CM on VLPSi-30.

**Figure 3.**
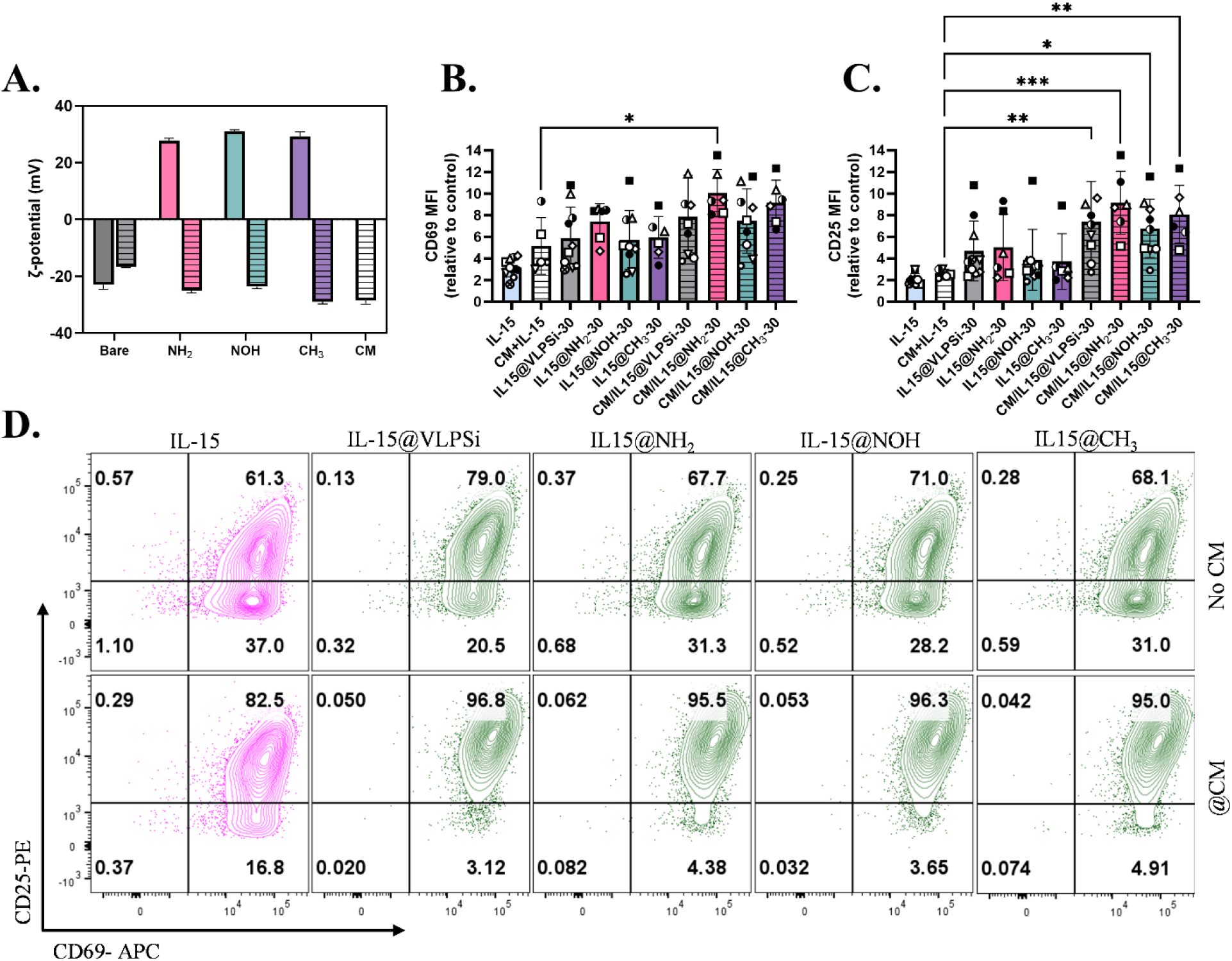
Amino modifications impact on NK cell activation. **A.** Zeta potential (ζ) of virus-like mesoporous silica nanoparticles (VLPSi) before and after surface modification: unmodified (grey), -NH_2_ (pink), NOH (green), and -CH_3_) (violet) modified, CM alone and CM-coated VLPSi shown with striped bars. **B-C**. Bar graphs showing normalized median fluorescence intensity (MFI) of **B**. CD69 and **C**. CD25 expression on NK cells. Peripheral blood mononuclear cells (PBMCs) were co-cultured with IL-15 coated virus-like mesoporous silica nanoparticles (IL15@VLPSis); grey) or IL-15@VLPSi with -NH_2_ (pink), -NHOH (green), -N(CH_3_)_3_) (purple) modifications, for 24-hours with and without MDA-MB-231 cell membrane (CM) coating (striped). IL-15 (blue) and CM alone (white) used as controls. **D**. Representative contour plot of CD25 and CD69 co-expression on NK cells. First row without CM coating and second row with CM coating (@CM). Data shown as mean ± SD. ***p < 0.001, **p < 0.01, *p < 0.05. Donors (n=7) are represented by different symbols.

The functionalized VLPSi-30 were evaluated for their ability to activate NK cells within PBMC from multiple donors. Modified VLPSi-30 were subsequently coated with IL-15 (IL15@VLPSi-30) and CM (CM/IL15@VLPSi-30) to assess combinatorial effects of cytokine delivery and CM coating. NK cell response varied between donors, reflecting the inherent heterogeneity of primary immune cells. Despite this variability, CM coating consistently increased the expression of CD25 (Fig. 3C, supplementary figure S4) as well as the CD25^+^CD69^+^ NK cell population compared to controls (IL-15 groups with or without CM; Figure 3D). Among the tested surface modifications, NH_2_-functionalized VLPSi-30 induced the strongest upregulation of both CD25 and CD69, particularly when combined with CM coating (IL15@NH_2_-30; Figure 3B and C). These findings indicate that VLPSi surface chemistry can be fine-tuned to improve NK cell activation, keeping in mind that the strongest effects are achieved through combination with CM coating. Based on these results, NH_2_ modified VLPSi-30 (NH_2_-30) were selected for the subsequent studies.

Mechanistic studies revealed that NK cell activation was driven primarily by CD14^+^ monocyte-NK cell crosstalk rather than direct VLPSi-NK cell interactions. Within PBMC, monocytes showed high uptake compared to minimal uptake by NK cells (Figure 1B), indicating that monocytes act as the primary sensors of VLPSi. Robust NK cell activation was observed within PBMC co-cultures, where γCM coating significantly upregulated CD25 and CD69 expression and IL15@NH_2_-30 showed significant upregulation in CD69 expression even without CM coating when compared to IL-15 control (IL-15) (Figure 4A). Consistently, depletion of CD14+ cells (Supplementary Figure S5) markedly reduced NK cell activation, evidenced by lower CD69 and CD25 expression (Figure 4B).

**Figure 4.**
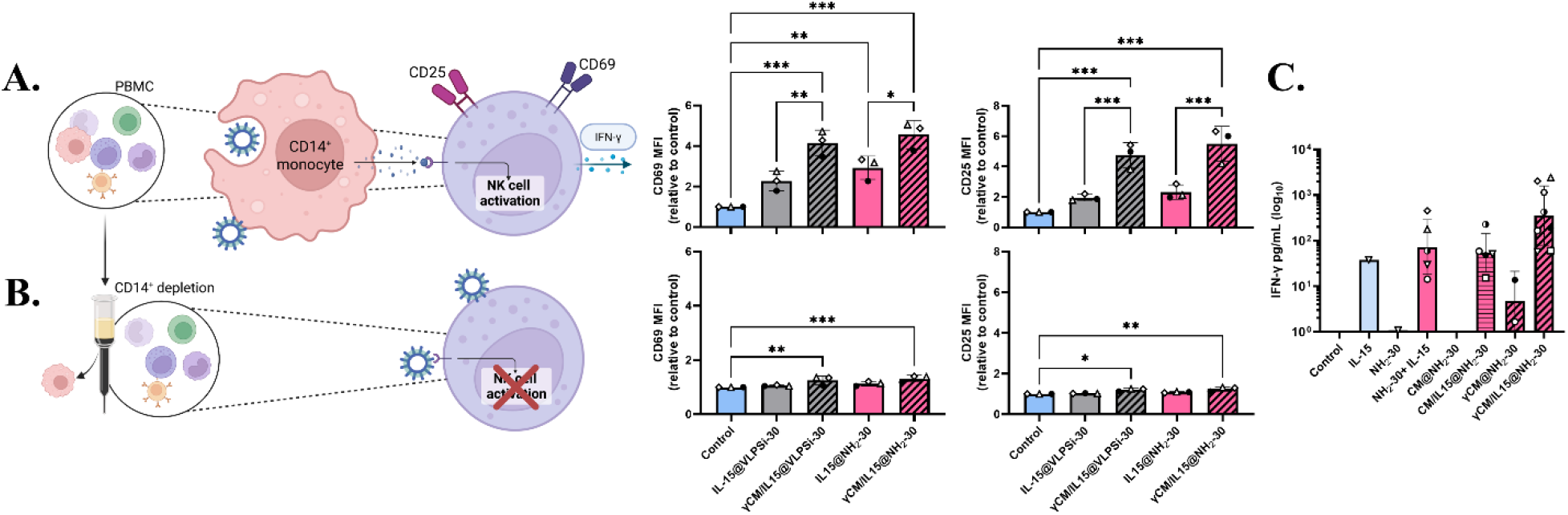
Impact from CD14-depletion. Peripheral blood mononuclear cells (PBMCs) and CD14-depleted PBMC were co-cultured with virus-like mesoporous silica nanoparticles (VLPSi) incorporated with IL-15 (IL15@VLPSi-30; grey) were compared to -NH_2_ functionalized VLPSi (IL15@NH2-30; pink) for 24-hours with and without IFN-γ treated MDA-MB-231 (MDA) cell membrane (γCM) coating (diagonal pattern). **A-B.** Bar graphs showing CD69 and CD25 expression by normalized median fluorescence intensity (MFI) on NK cells in peripheral mononuclear cells (PBMCs) **A**. before and **B**. after CD14-depletion. **C**. Bar graph showing log_10_-transformed IFN-γ levels (pg/mL;) measured from PBMCs co-cultured with NH_2_-30 VLPSi with and without IL-15 and CM coating. CMs were derived from MDA (CM; stripes) and γCM (diagonal pattern). Data shown as mean ± SD. ***p < 0.001, **p < 0.01, *p < 0.05. Donors represented by different symbols.

Monocyte-dependent activation was further supported by IFN-γ secretion, which was significantly elevated in PBMC co-cultures in the presence of IL-15, and under detection limit after CD14^+^-depletion. The highest IFN-γ levels were observed with IL15@NH_2_-30, particularly when combined with γCM coating (diagonal stripes; Fig. 4C).

These results indicate that downstream NK cell responses arise indirectly through monocyte-mediated signalling rather than direct contact with VLPSi. This mechanism aligns with a recent proteomics-guided mechanistic study demonstrating spike-length-dependent activation of TLR, NF-κB, MAPK, RIG-I receptor pathways in murine macrophages, with upregulation of TRAF6 and ERK1/2 expression and elevated TNF-α/IL-1β transcription, supporting myeloid PRR engagement by virus-like topology [19]. Accordingly, previous studies have shown that depletion of CD14^+^ cells reduces NK cell IFN-γ secretion due to loss of monocyte-derived cytokine signalling [29], supporting our observation that monocytes are critical for NK cell activation. Together with our findings, monocyte activation is positioned as the primary trigger of VLPSi-mediated NK stimulation in PBMCs. These findings are consistent with the known immune cell biology: monocytes function as professional phagocytes and regulators of innate immune communication [30], whereas NK cells typically engage their targets through transient immune synapses rather than endocytic pathways [31].

Engineered K562 cells are widely used in feeder-cell based NK cells expansion [32] and have been used to produce nanoscale plasma membrane vesicles, which have been shown to activate NK cells *ex vivo*, and *in vivo* [33]. These cells present membrane-bound activating ligands capable of directly engaging NK cell receptors. We hypothesized that incorporating CM derived from K562 feeder cells onto VLPSi could enhance NK receptor engagement, potentially strengthening their activation and *ex vivo* expansion even in the absence of accessory monocytes. Purified naïve NK cells were co-cultured with CM-coated IL15@NH_2_-30 VLPSi. CMs were derived from γCM and genetically engineered K562 feeder cell variants: unmodified K562 cells (K562A), cells engineered to express membrane-bound IL-15 and 4-1BB ligand (K562B), and membrane-bound IL-21, CD48, and 4-1BB ligand (K562C), commonly used in NK cell *ex vivo* expansion [23]. Naïve NK cell activation remained low with all, except K562C-CM coating, which significantly increased the CD69 expression (Figure 5A) in all donors. However, robust IFN-γ secretion and CD25 upregulation still required the presence of CD14^+^ monocytes.

**Figure 5.**
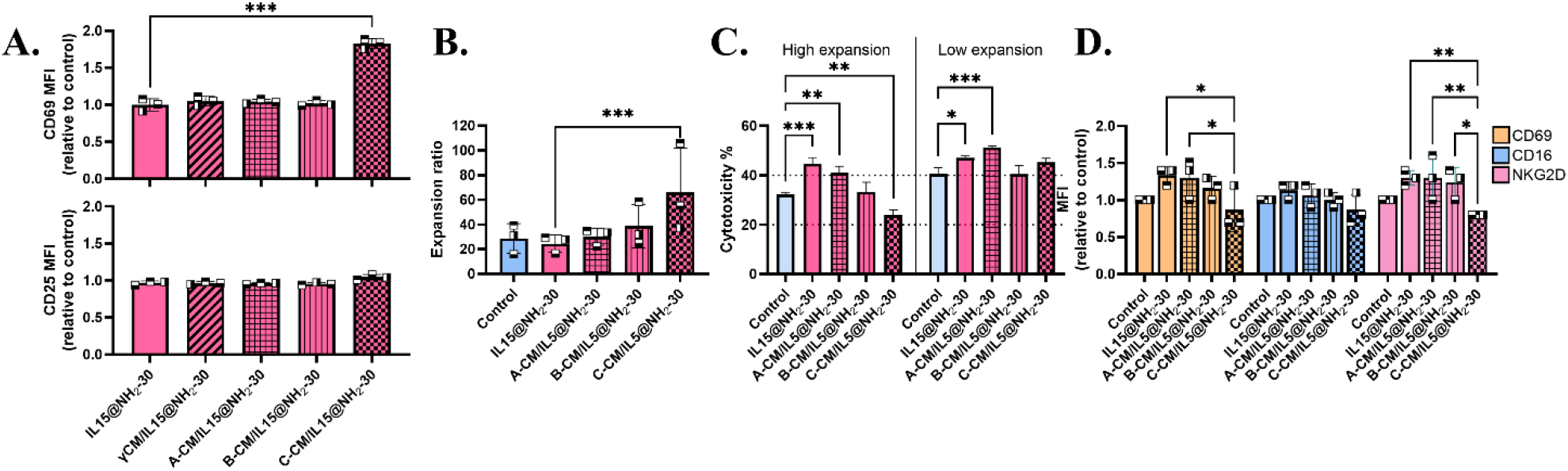
Cell membrane (CM)-coated virus-like mesoporous silica nanoparticles (VLPS NPs) impact NK cell expansion. **A.** Naïve NK cells were activated NH_2_-modified VLPSi incorporating IL-15 (IL15@NH_2_-30) with and without CM coating. CMs were derived from unmodified (A; grid), and genetically modified K562 cells expressing membrane-bound IL-15 and 4-1BB ligand (B; stripes), and membrane-bound IL-21, CD48, and 4-1BB ligand (C; check). Bar graphs show CD69 and CD25 expression by normalized median fluorescence intensity (MFI) naïve purified NK cells after 24 hours co-culturing. **B**. Bar graph showing NK cell expansion ratio. Naïve NK cells were expanded for 17 days in media supplemented with IL-2 and IL-15 and activated on days 3 and 10 with IL15@NH_2-_30, A-CM/IL15@NH_2_-30, B-CM/IL15@NH_2_-30 or C-CM/IL15@NH_2_-30. **C**. Bar graphs show the NK cell cytotoxicity of highest and lowest expansions with CM-coated VLPSi. MDA-MB-231 cells were co-cultured with NK cells (10:1 E:T ratio) for 24 hours. **D**. Bar graphs showing normalized median fluorescence intensity (MFI) expression of CD69, CD16 and NKG2D expression on NK cells after expansion. Data shown as mean ± SD. ***p < 0.001, **p < 0.01, *p < 0.05. Donors represented by different symbols, n= 3.

K562 CM coating showed promise in NK cell *ex vivo* expansion studies. CM coating could introduce immune co-stimulatory mimicking ligands, without the need for intact feeder cells and could provide a safety advantage for NK cell expansion [34]. Used CMs were derived from K562A (A-CM), K562B (B-CM) and K562C (C-CM) and coated to IL-15@NH_2_-30. Naïve NK cells were cultured with well-established NK cell expansion protocol, in which IL-15 and IL-2 were included during expansion, with feeder cells replaced by CM-coated VLPSi. NK cells were expanded for 17 days, with VLPSi added on days 3 and 10, after which the yield, phenotype and functionality were assessed. NK cell yield reflected the results of the short-term activation assay (Fig. 5A). K562C-CM-coated VLPSi (C-CM/IL15@ NH_2_-30) induced the highest fold expansion (66±36), exceeding expansion rates from cytokine only control (29±7), bare VLPSi (25±7; IL15@NH_2_-30) and K562A (30±7; A-CM/IL15@NH_2_-30) or K562B (39±17; B-CM/IL15@NH_2_-30) CM-coated VLPSi (Fig. 5B). These results indicate that membrane composition influenced NK cell proliferation, consistent with the presence of activating ligands (IL-21, CD48, and 4-1BB) in K562C-derived CM. This expansion approach could be further developed by incorporating additional monocyte-derived signals, such as IL-18, onto the VLPSi surface, which have been shown advantage in NK cell expansions [36,37].

NK cell activation markers (CD69, CD25, CD16 and NKG2D) were assessed following expansion. CD25 expression was downregulated in all conditions (data not shown), which is a known effect from the use of IL-2 during NK cell expansion [21]. CD69 and NKG2D expression were significantly lower in NK cells expanded with K562C-CM-coated VLPSi (Fig. 5C; C-CM/IL15@NH_2_-30), which also induced the highest proliferation rates. This phenotype suggests a shift toward an expansion-dominant state, aligning with previous reports that strong proliferative stimuli can differentially modulate activating receptor expression during *ex vivo* expansion [21,24].

To determine whether these phenotypic changes were associated with functional alterations, NK cell cytotoxicity was evaluated in donors representing the highest and lowest expansion levels with C-CM/IL15@NH_2_-30 against MDA cells at 10:1 E:T ratio. NK cells exhibiting the highest expansion showed reduced cytotoxicity, whereas NK cells conditions associated with lower expansion had higher cytotoxicity (Fig. 5C). Uncoated VLPSi (IL15@NH_2_-30) and K562A-CM-coated VLPSi (A-CM/IL15@NH_2_-30) expansions increased CD69 and NKG2D expression (Fig. 5D) and enhanced the NK cell cytotoxicity (Fig. 5C) compared to the cytokine control. This was seen with both donors. These findings suggest that distinct CM compositions could be utilized to differentially modulate NK cell activation and function *ex vivo*.

This inverse relationship between expansion, activation marker expression (CD69, NKG2D), and cytotoxic function is consistent with previously reported trade-offs in NK cell biology, where strong proliferative stimuli can drive cells toward an expansion-dominant but functionally attenuated phenotype. Together, these findings demonstrate that CM-coated VLPSi enable NK cell expansion and modulate effector function in a CM-dependent manner, providing a strategy to optimize NK cell products for therapeutic applications.

The results established CM-coated VLPSi as a tunable platform in which virus-like topology, surface chemistry, cytokine presentation, and membrane composition together can be utilized to shape the immune response. CM derived from different origins could be used as a powerful tool for incorporating specific immunomodulatory signals into the nanomaterial design. This work provides a platform for nanomaterial design able to engage innate immune crosstalk to modulate NK cell activity and hold potential for NK cell-based cancer immunotherapies.

## Acknowledgement

This work was supported by the Finnish Cancer Foundation (Grant no. 230130) and Research Council of Finland (Grant no. 356056). The authors thank Satu Kaipainen for valuable support and technical assistance during blood sampling. We also acknowledge technical support from Petri Mäkinen and Tiina Nieminen (A.I Virtanen Institute for Molecular Sciences), and UEF Cell and the Tissue Imaging Unit, and SIB Labs.

