## Supporting information for "Biomimetic virus-like mesoporous silica nanoparticles activate NK cells indirectly via monocyte crosstalk"

[3] iCell, Research and Development, Finnish Red Cross Blood Service, Haartmaninkatu 8, 00290 Helsinki, Finland

### Supplementary Methods

#### 1.1 Synthesis and surface functionalization of VLPSi

VLPSi ynthesis was adapted from [1]. Briefly, hexadecyltrimethylammonium bromide (CTAB; Sigma) was dissolved in deionized water to a final concentration of 1%. The mixture was stirred at 60 °C until fully dissolved, after which 0.1 M sodium hydroxide (NaOH; Fisher chemical) was added. When the mixture was homogenous, a 1:4 mixture of tetraethyl orthosilicate (TEOS; Merck) and cyclohexane (Merck) was added using a syringe pump. The reaction time for preparing VLPSi with 5 nm and 30 nm spikes was 48 and 70 hours, respectively. After the reaction, the VLPSi were washed twice by centrifugation (12 700 g, 10 min) with ethanol, followed by calcination in a tube furnace at 600 °C in air for 4 h.

VLPSi were functionalized post-synthesis using silane coupling agents. 3-Aminopropyltriethoxysilane (APTES; 99% purity; VWR), N,N-BIS(2-hydroxyethyl)-3-aminopropyltriethoxysilane (SIB1140.0; 62% in Ethanol; AcroSea), and N-[3-(Trimethoxysilyl)propyl]-N,N,N-trimethylammonium chloride (50% in Methanol; Thermo Scientific) were used to prepare VLPSi-$NH_2$, VLPSi-NHOH, and VLPSi-$N(CH_3)_3$, respectively. Functionalization was done by stirring at 65 °C in ethanol for 1 hour. After the reaction, the product was washed twice by centrifugation (12 700g, 10 min) with ethanol. The final VLPSi were stored in ethanol at room temperature (RT) for further use.

### 1.2 Cell culture

MDA-MB-231 (MD Anderson-metastatic breast-231; breast cancer) were cultured in RPMI-1640 (VWR), and MCF-7 (Michigan Cancer Foundation-7; breast cancer) in DMEM (VWR), both media were supplemented with 10% fetal bovine serum (FBS, gamma irradiated, Gibco) and 100 U/ml penicillin and 100 µg/ml streptomycin (P/S, Gibco). Cells were maintained in a humidified atmosphere at 37 °C, with 5% $CO_2$ and passaged when reaching 80-90% confluency. Cells were tested for mycoplasma with GenieColor Mycoplasma Detection Kit (AssayGenie). K562 cells were received cryopreserved from the iCell Group (as described in Saari et al. submitted).

### 1.3 Cell membrane extraction

Cell membranes (CMs) were extracted from MDA-MB-231 and IFN-γ (50 ng/mL) treated MDA-MB-231 cells, and three genetically modified K562 cell lines: K562 (K562A), K562-mbIL15-41BBL (K562B), and K562-mbIL21-CD48-41BBL (K562C), using our previously described method [2]. Briefly, 50-100x$10^6$ cells were washed three times with PBS, suspended to cold hypotonic lysing buffer [20 mM Tris-HCl pH 7.5 + 10 nm KCl + 2 mM $MgCl_2$; supplemented with EDTA-free mini protease inhibitor tablet (1 x 10 mL solution; Roche)] and lysed on rotation at 4°C for 30 min. Cells were then mechanically disrupted using Dounce homogenizer, subjecting them twice to 100 passes. Supernatants were collected after each step and purified by centrifugation, first 3200 g 5 min and then 10 000 g for 10 min, pellets were discarded after each step. CMs were concentrated with two step ultracentrifugation (100 000 g for 1 hour followed by 100 000 g for 30 min), pellet was washed in between with HEPES buffer (pH 7.5) and stored at -80°C in HEPES. Protein concentration was determined using bicinchoninic acid (BCA) assay (Pierce™ Dilution-Free™ Rapid Gold BCA Protein Assay Kit, Thermo Fisher) according to manufacturer´s instructions.

### 1.4 Cell membrane coating

VLPSi were coated with CM at a 35% (w/w) CM to VLPSi ratio in 50% PBS. The ratio was optimized by quantifying CM bound to VLPSi after removing unbound CM by centrifugation (12 000 g, 10 min), showing that 80-99% of the added membrane was bound to the VLPSi (Supplementary Figure S1). CM coating was done using an Elmasonic S10 H ultrasonic bath (Singen; 5 min on ice) and confirmed by changes in zeta potential (ζ; see below) and by protein concentration (BCA assay). The CM-coated VLPSi (CM@VLPSi) were further suspended in cell culture media for the activation assays.

#### 1.5 VLPSi characterization

The VLPSi morphology and spike length were characterized by transmission electron microscopy (TEM). Bare and CM-coated were plated onto formwar and carbon coated glow discharged copper grids (200 mesh; Electron Microscopy Sciences). Grids were protected from light and exposed to osmium tetroxide ($OsO_4$; Electron Microscopy Sciences) vapor for 10 minutes. Samples were imaged with JEM-2100F (JEM Ltd., Japan).

Zeta potential (ζ) was measured using a Malvern Zetasizer (Malvern Zetasizer Nano ZS). Samples were dispersed in distilled water and measured in folded capillary cells (Fisher Scientific, NC0491866). An average of three measurements (60 s) were recorded per sample. Colloidal stability was measured, in PBS, on the same instrument at four timepoints (1, 4, 8 and 24 hours) by dynamic light scattering (DLS) using disposable cuvettes and the volume-weighted size distribution mode under identical dispersion conditions. An average of three measurements (60 s) were recorded per sample.

#### 1.6 SDS-Page and Western Blotting

CM coating was further analyzed using SDS-PAGE and Western blot analysis. 25 μg of protein per sample was suspended in 4× Laemmli sample buffer and denatured at 95 °C. Proteins were separated using 10% sodium dodecyl sulfate–polyacrylamide gel electrophoresis (SDS-PAGE) gel and immobilized onto nitrocellulose-membrane (Thermo Fisher Scientific). After washing, the membrane was stained with Ponceau S staining solution (5 min at RT) with agitation. Excess stain was washed until protein bands were clearly visible. For Western Blotting the membrane was washed with Tris-buffered saline with Tween-20 (TBST) before blocking with 3% bovine serum albumin (BSA; Thermo Scientific) in PBS for 30 minutes. Membrane was stained with primary antibodies [rabbit polyclonal anti-HLA-F (1:1000; Proteintech, Cat. No. 14670-1-AP) and rabbit polyclonal anti-Lamin B1 (1:5000; Abcam, Cat. No. Ab16048, Cambridge, UK)] and subsequently incubated with HRP-conjugated goat anti-rabbit IgG secondary antibody (Invitrogen, Thermo Fisher Scientific). Chemiluminescent signal was developed using Clarity Western ECL Substrate (Bio-Rad Laboratories) and measured with ChemiDoc MP Imaging System (Bio-Rad Laboratories).

### 1.7 Confocal Microscopy

Co-localization of VLPSi and CM was assessed to verify uptake of CM-coated VLPSi. 1 x $10^5$ THP-1 cells per well were seeded into an 8 well ibidi plate (ibidi, Germany) and allowed to attach overnight at 37°C, 5% $CO_2$. CM was stained with DiI (1,1′-dioctadecyl-3,3,3′,3′-tetramethylindocarbocyanine perchlorate; Sigma-Alrich) in HEPES (1:1000) overnight on rotation at 37°C. The stained CM was washed and collected using Optima™ XE-100 Ultracentrifuge (100 000 g 30 min; Beckman Coulter). VLPSi-$NH_2$ were labeled with Fluorescein 5(6)-isothiocyanate (FITC, Alfa Chemical; 1 mg/mL) overnight on rotation at RT, and washed by centrifugation (14 000 g, 10 min) with ethanol. CM coating was done as described above. Target cells were co-cultured with labelled VLPSi and CM@VLPSi for 4 hours. Nuclei were stained with DAPI (Sigma-Alrich) and fluorescent images were obtained with a Zeiss Axio Observer inverted microscope (40×oil-objective) with LSM800 confocal module (Carl Zeiss Microimaging GmbH, Jena, Germany).

### 1.8 Internalization

Target cells (PBMC, expanded NK cells (exNK), MDA and THP-1) were co-cultured with the FITC labelled VLPSi (VLPSi-FITC; see above) and CM-coated VLPSi-FITC for 4 hours. Cells were stained with 7-AAD, CD56, CD3, and CD14 antibodies and analyzed using flow cytometry (see below). FITC positive cells were gated and analyzed from live single-cell population after excluding cell debris and doublets. Cell type specific gating was used for PBMC [NK cells ($CD56^+$/$CD3^-$) and monocytes ($CD14^+$)].

### 1.9 Biocompatibility

MDA-MB-231 and MCF-7 cells were seeded into 96-well plates, 1x$10^4$ cells per well, and allowed to attach overnight. VLPSi and CM@VLPSi were added as concentration series (0, 13, 25, 50, 100, and 200 µg/mL), and cell viability was measured after 24 hour co-culturing using CellTiter-Glo®2.0 assay (Promega) according to manufacturer's instructions. Luminescence was measured using VICTOR Nivo™ microplate reader (Revvity). After blank subtraction the cell viability was calculated relative to the control.

#### 1.10 PBMC isolation

Human peripheral blood mononuclear cells (PBMCs) were isolated from peripheral blood samples diluted 1:1 with PBS by density gradient centrifugation using Histopaque®-1077 HybriMax™ and SepMate™ -50 tubes (StemCell) according to manufacturer's instructions. PBMCs were cryopreserved in FBS (Sigma) supplemented with 10% DMSO (Sigma).

#### 1.11 CD14 depletion

$CD14^+$ cells were depleted from PBMCs using CD14 MicroBeads and LD columns according to the manufacturer's instructions. MACS® MultiStand and QuadroMACS™ Separator (Miltenyi Biotec) were used for magnetic separation.

#### 1.12 NK cell activation studies

PBMCs and NK cells (expanded or naïve) were seeded to 24-well plate at 0.5 x $10^6$ cells per well in culture medium (RPMI-1640 + 10% FBS + P/S) and co-cultured with 50 µg/mL VLPSi or CM@VLPSi for 24 hours. The used time point and concentration were selected based on preliminary studies (Supplementary Figure S2). Cells were analyzed using flow cytometry, and the co-culture medium was used for IFN-γ assays.

Human IFN-γ release was quantified using the Lumit® IFN-γ (Human) Immunoassay (Promega, TM686). Media samples were frozen and thawed prior to analysis. The assay was performed according to the manufacturer's instructions. Briefly, luminescence generated by NanoBiT®-based detection reagents upon binding to human IFN-γ was measured using a VICTOR Nivo™ microplate reader (Revvity).

#### 1.13 Flow cytometric characterization

Staining protocol was adapted from [3]. Staining was done in flow buffer [phosphate buffered saline (PBS, Gibco) + 0.02% Sodium Azide ($NaN_3$, Sigma) + 2 mM ethylenediaminetetraacetic acid (EDTA, Sigma) + 2% FBS (Gibco)]. Cells were blocked with 10% human FcR Blocking Reagent (Miltenyi Biotec) in flow buffer, followed by staining for 15 min at RT with anti-human monoclonal antibodies (mAb). The used antibodies, indicated as target, fluorochrome and clone, were as follows: CD3-BV510 (OKT3), and CD56-BV421 (5.1H11) for identification of NK cells (CD3-/CD56+); CD25-PE (M-

A251), CD16-FITC (3G8), CD69-APC (FN50), and NKG2D-APC-Fire750 (1D11) as NK activation markers; CD14-APC-Fire750 (1D11) as monocyte marker. All mAbs were obtained from Biolegend, except 7-amino-actinomycin D (7-AAD; Miltenyi). Cells were acquired on a CytoFLEX S (Beckman Coulter) and analyzed using CytExpert v2 (Beckman Coulter) and FlowJo v10 (BD Life Sciences, TreeStar). All samples were gated using same strategy (Supplementary Figure 3A). Briefly, dead cells and debris were excluded based on forward and side scatter, and live cells were identified from single cells (FSC-A vs. FSC-H) by 7-AAD expression. NK cells ($CD56^+/CD3^-$) cells were subsequently analyzed for expression of CD69 and CD25. Mean fluorescence intensity (MFI) values were normalized to the control condition for each donor to account for donor variability and to assess changes in activation.

#### 1.14 NK cell isolation and expansion

Human PBMCs were separated from buffy coats by density gradient centrifugation using Ficoll-Paque (GE Healthcare) and NK cells ($CD56^+/CD3^-$) were isolated using the NK Cell Isolation Kit (Miltenyi Biotec) by negative selection according to the manufacturer's instructions. Naïve NK cells were expanded for 17 days (to expanded NK cells, exNKs) in NK expansion medium [NK MACS medium (Miltenyi Biotech) + 1% NK MACS supplement (Miltenyi Biotech) + 5% Human AB Serum (Sigma) + 500 IU/mL human IL-2 (Miltenyi Biotech) + 140 IU/ml human IL-15 (Miltenyi Biotech) + 1% penicillin/streptomycin (Gibco)]. On days 3 and 10, cells were stimulated with irradiated K562-mbIL21-CD48-41BBL (K562C) feeder cells at a 1:5 NK to feeder cell ratio. On day 17, NK cells were collected and cryopreserved.

CM@VLPSi-expanded NK cells were activated and maintained as described above. On days 3 and 10, cells were stimulated with 50 µg/mL CM@VLPSi. CMs were derived from three different feeder cell lines (K562A, K562B, and K562C; see above). Cell count and viability were determined using NucleoCounter® NC 202™ automated cell counter (ChemoMetec).

#### 1.15 NK cell cytotoxicity assay

Target cells (MDA-MB-231 and MCF-7) were seeded into flat 96-well plates at $1 \times 10^4$ cells per well and allowed to attach at 37°C in 5% $CO_2$ overnight. Bare and CM-coated VLPSi were added at 50 µg/mL concentration, together with and NK cells (effectors) at three effectors to target (E:T) ratios (10:1, 5:1, 2.5:1). After 24 hour co-culturing the cytotoxicity was measured using CellTiter-Glo®2.0

assay (Promega). Luminescence was measured using VICTOR Nivo™ microplate reader (Revvity). The NK cell cytotoxicity was calculated according to the following formula:

$$Cytotoxicity\% = \left(1 - \frac{RLUsample - RLUeffector}{RLUtarget}\right) x\ 100\% \quad | RLU = Luminescence$$

#### 1.16 Statistical analysis

Statistical analyses were performed using GraphPad Prism 10. Comparisons between groups were performed using two-way ANOVA with Tukey's or Dunnett's post hoc test. $P < 0.05$ was considered statistically significant. Results are shown as mean ± SD.

#### 1.17 Ethics

Peripheral blood samples were collected from voluntary blood donors with informed consent and handled under an approval by the Ethics Committee of the Hospital District of Northern Savo (295/13.00/2024). Primary NK cells were obtained from buffy coats, which were residual products from blood donations (Blood Component Delivery License Decision 178/7/2024/iCell group).

#### CRediT authorship contribution statement

**Minna Sivonen**: Conceptualization, Methodology, Investigation, Data curation, Visualization, Writing - original draft, **Silja Saarela**: Investigation, Methodology, **Jiajia Wang**: Investigation, Writing - review & editing, **Mira Saari**: Investigation, Methodology, Writing - review & editing, **Emmi Järvelä**: Investigation, Writing - review & editing, **Lotta Andersson**: Investigation, **Enkhzaya Batnasan**: Investigation, Methodology, Writing - review & editing, **Leena Latonen**: Conceptualization, **Helka Göös**: Conceptualization, Writing – review & editing, **Vesa-Pekka Lehto:** Supervision, Writing - review & editing, **Wujun Xu:** Supervision**,** Conceptualization**,** Methodology**,** Writing - review & editing, Funding acquisition

#### Declaration of AI and AI-assisted technologies in the writing process

During the preparation of this work the author(s) used Microsoft co-pilot in order to improve readability and language of the publication. After using this tool/service, the author(s) reviewed and edited the content as needed and take(s) full responsibility for the content of the publication.

### Supplementary Figures

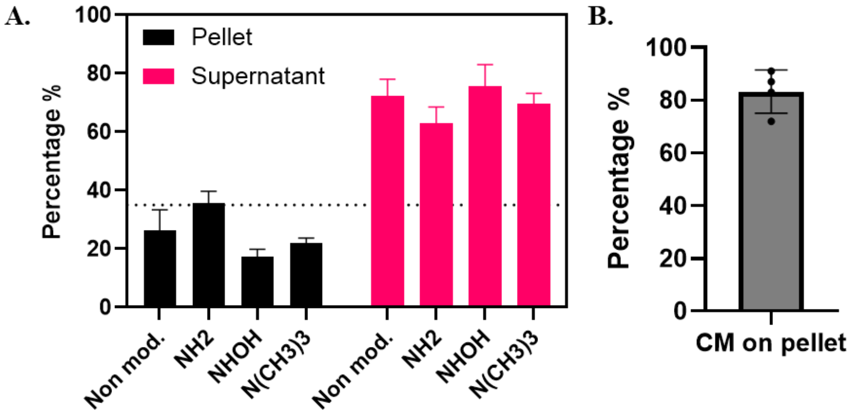


**Supplementary Figure S1.** Coating optimization studies. **A.** Bar graph showing coating efficiency from pellet and supernatant following 1:1 (w/w) coating of VLPSi with cell membrane (CM). **B.** Bar graph showing 4 separate coating studies done with 0.35:1 (35%) coating. Average coating efficiency 82%.

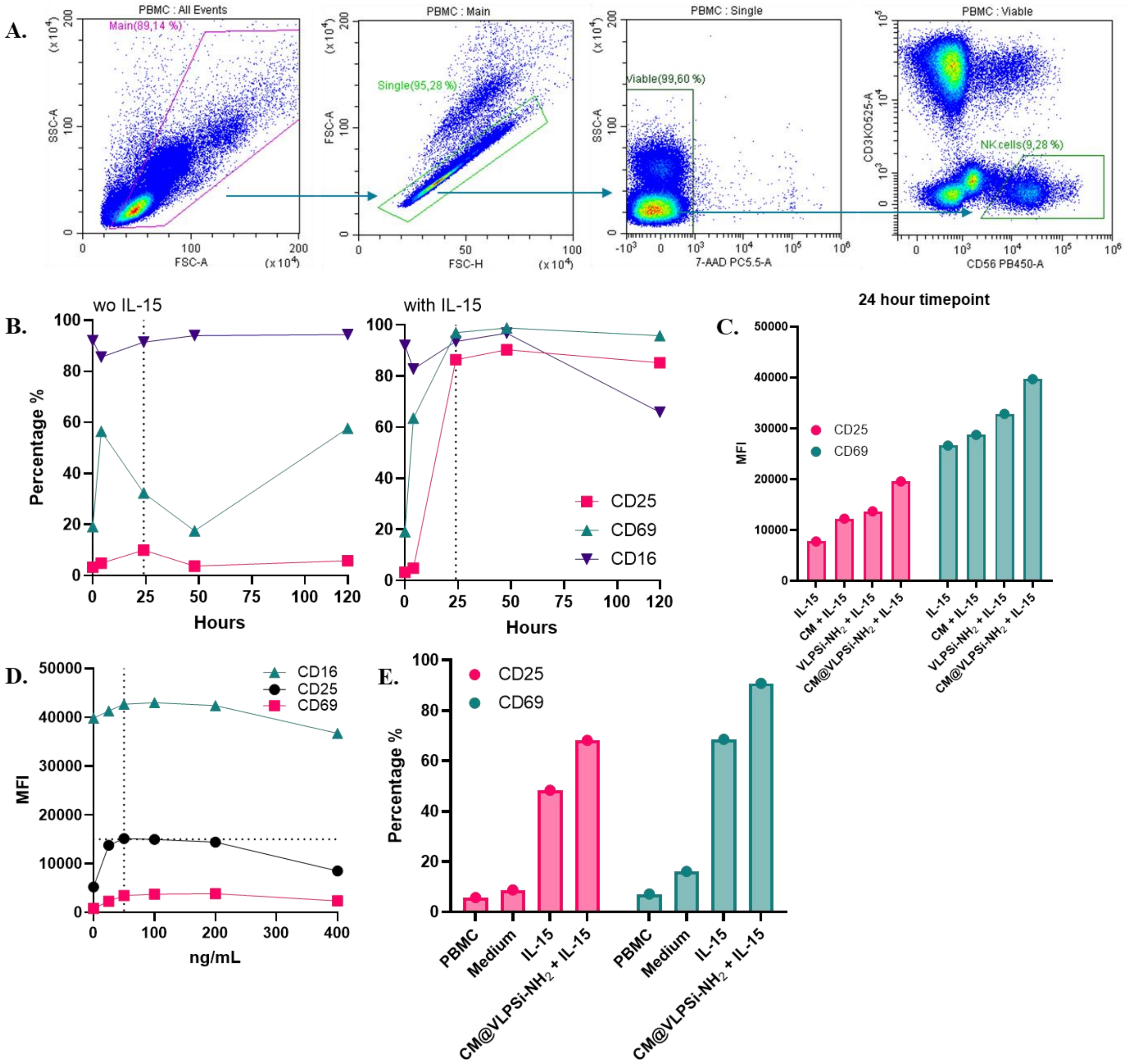


**Supplementary Figure S2. A.** Representative example for the gating strategy for viable (7-AAD$^-$) CD3$^-$/CD56$^+$ NK cells. **B.** Timepoint study for NK cell activation. Line graph shows percentage of CD25 (pink), CD69 (green) and CD16 (violet) expressing NK cells after 4 hours, 24 hours, 48 hours and 5 days co-culturing with amino modified MDA-MB-231cell membrane (CM)-coated VLPSi-30 (CM@VLPSi-$NH_2$). Results with and without supplementary IL-15 (10 ng/mL). **C.** Bar graph showing CD25 and CD69 median fluorescence intensity (MFI) after 24 hour co-culturing with IL-15 supplemented CM, VLPSi-$NH_2$, and CM@VLPSi-$NH_2$. **D.** Concentration study with CM@VLPSi-$NH_2$, with IL-15, at 24 hour timepoint. Line graph showing concentration dependent change in CD16, CD25 and CD69 MFI intensity. E. Bar graph showing CD25 and CD69 median fluorescence intensity (MFI) after 24 hour co-culturing with 50 ng/mL VLPSi-$NH_2$, and CM@VLPSi-$NH_2$ (supplemented with IL-15).

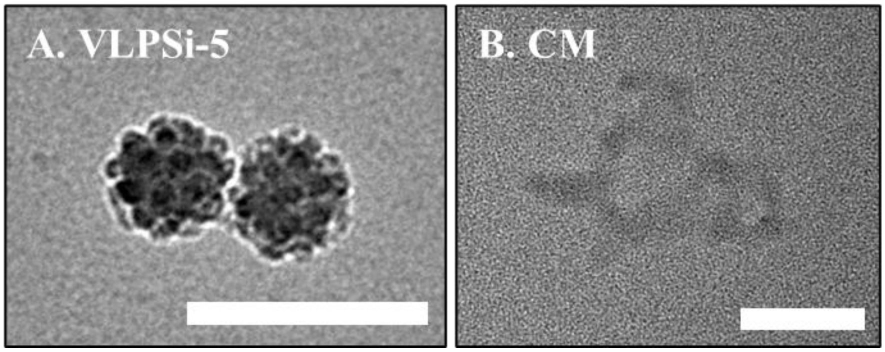


**Supplementary Figure S3.** Transmission electron microscope (TEM) images from **A.** VLPSi with 5 nm spikes (VLPSi-5), and **B.** Free cell membrane (CM)

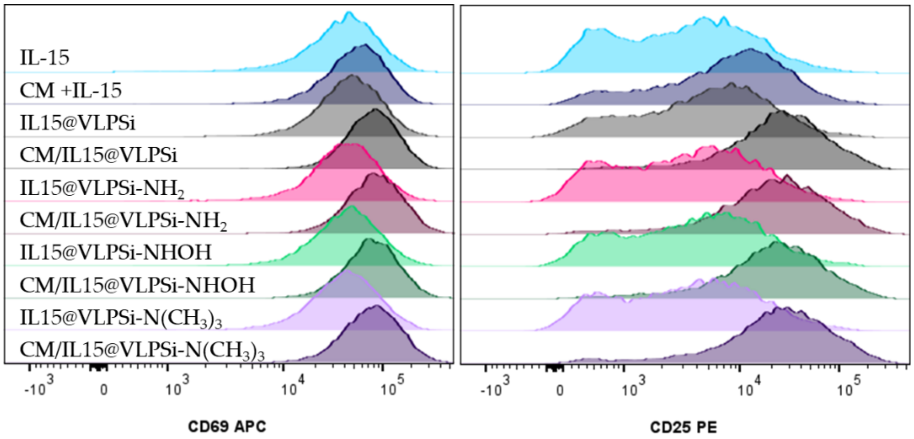


**Supplementary Figure S4.** Representative histogram of CD69 and CD25 expression after 24 hour co-culture with differently amino modified VLPSi, with and without cell membrane (CM) coating.

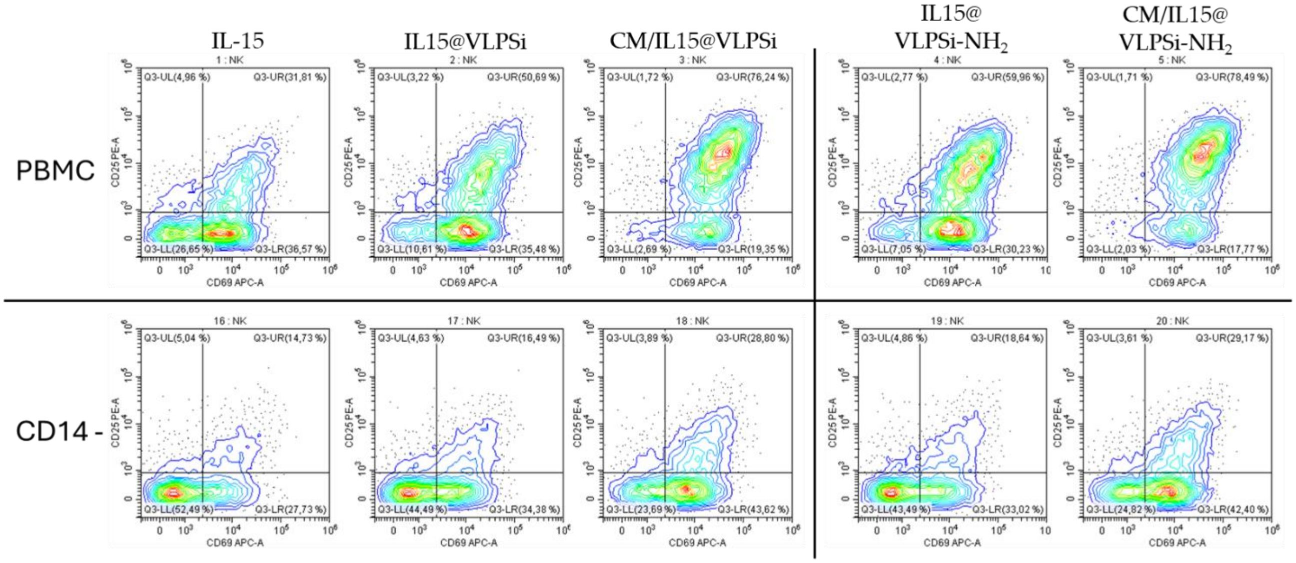


**Supplementary Figure S5.** Representative co-expression plots of CD25 and CD69 on NK cells after 24 hour co-culture with VLPSi and VLPSi-NH2 with and without MDA-MB-21 cell membrane (CM) coating. First row shows the expression on PBMC-co-cultures and second row after CD14 depletion.
